# BaRDIC: robust peak calling for RNA-DNA interaction data

**DOI:** 10.1101/2023.09.21.558815

**Authors:** Dmitry E. Mylarshchikov, Arina I. Nikolskaya, Olesja D. Bogomaz, Anastasia A. Zharikova, Andrey A. Mironov

## Abstract

Chromatin-associated non-coding RNAs play important roles in various cellular processes by targeting genomic loci. Two types of genome-wide NGS experiments exist to detect such targets: “one-to-all”, which focuses on targets of a single RNA, and “all-to-all”, which captures targets of all RNAs in a sample. As with many NGS experiments, they are prone to biases and noise, so it becomes essential to detect “peaks” – specific interactions of an RNA with genomic targets. Here we present BaRDIC – Binomial RNA-DNA Interaction Caller – a tailored method to detect peaks in both types of RNA-DNA interaction data. BaRDIC is the first tool to simultaneously take into account the two most prominent biases in the data: chromatin heterogeneity and distance-dependent decay of interaction frequency. Since RNAs differ in their interaction preferences, BaRDIC adapts peak sizes according to the abundances and contact patterns of individual RNAs. These features enable BaRDIC to make more robust predictions than currently applied peak-calling algorithms and better handle the characteristic sparsity of all-to-all data. BaRDIC package is freely available at https://github.com/dmitrymyl/BaRDIC.

## INTRODUCTION

Chromatin-associated non-coding RNAs play important roles in various cellular processes (1, 2, 3). Many sequencing techniques are now available to localize RNAs on the whole chromatin, which can be broadly classified into “one-to-all” (OTA) and “all-to-all” (ATA) methods (4, 5). OTA methods (6, 7, 8, 9) hybridize biotinylated RNA-specific oligonucleotide probes to find the contacts of a single target RNA. High-throughput ATA methods (10, 11, 12, 13, 14, 15) detect genome-wide contacts of all potential chromatin-associated RNAs using RNA-DNA proximity ligation (11, 16). Identifying specific interactions of an individual RNA with a particular genomic location, a process called “peak-calling”, has been a focus of all RNA-DNA interaction studies. However, RNA-DNA interaction data contain biological and technology-specific biases, which hinder the detection of real RNA-binding sites on the chromatin. Similar to peak-calling algorithms in ChIP-seq data and chromatin loop detection in Hi-C data, there is a need to develop tailored methods for identifying specific interactions in RNA-DNA interaction data.

Specific interaction loci can be defined as genomic regions where the number of contacts is significantly higher than expected in a given background model. Ideally, the background model captures all possible sources of biases. One of these biases is chromatin heterogeneity, which includes chromatin accessibility, amplification bias, and copy number variation. This bias is also present in ChIP-seq datasets. And since the nature of OTA data is also similar to that of ChIP-seq data due to common experimental steps, a ChIP-seq specific peak caller MACS2 is usually applied to OTA data, using an input track, which evaluates chromatin heterogeneity. For ATA data, chromatin heterogeneity can be inferred from the dataset itself, according to GRID data processing protocol (11). The endogenous background here is based on *trans* chromosomal interactions of mRNAs, assuming such contacts are mostly non-specific. In contrast, RADICL protocol (16) proposed a uniform binomial model that ignores the heterogeneous nature of the chromatin.

Another source of bias is the decay of contact density with increasing distance from the RNA transcription site. This effect was described in several studies dedicated to RNA-DNA interaction data analysis (15, 16). We observed this distance-dependent effect in both OTA and ATA datasets (Figure 1A) with our previously developed RNA-Chrom database (17). Analogous to Hi-C, we will refer to this phenomenon as “scaling” (18). MACS2, GRID, and RADICL approaches do not consider scaling in their models, therefore, these algorithms might overestimate the statistical significance of contacts neighboring an RNA source gene.

**Figure 1.**
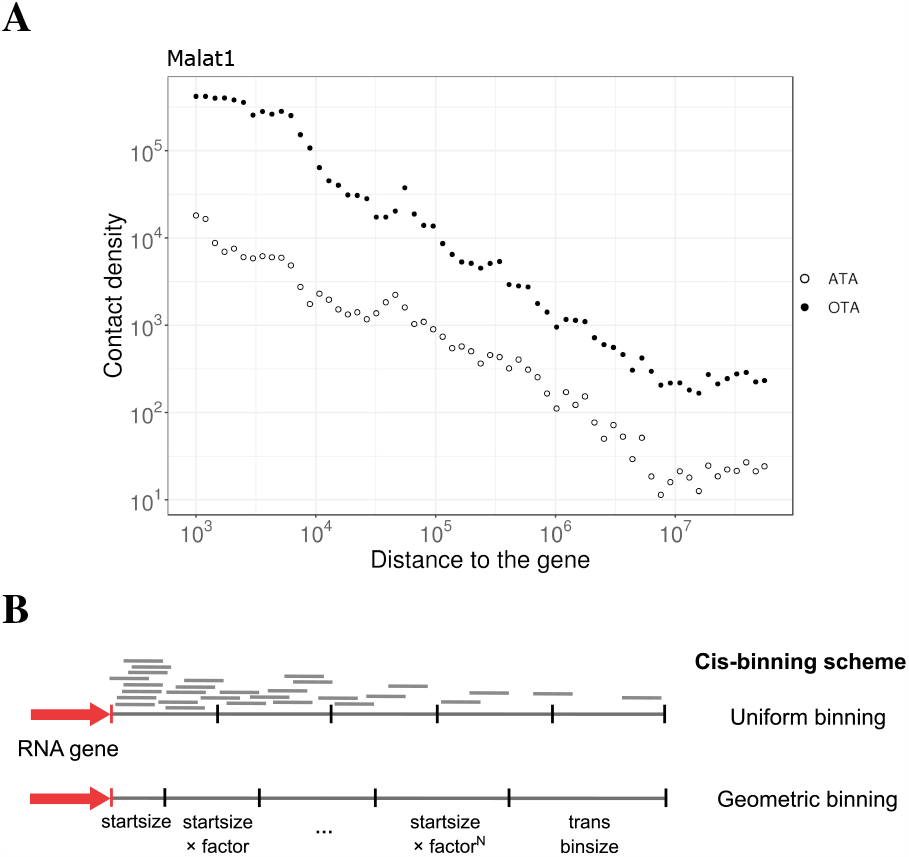
**(A)** Dependency of contact density on the distance between the RNA source gene and chromatin target loci (scaling) in double logarithmic coordinates for Malat1. Black points: ChIRP OTA data on mESC (19), white points: ATA GRID data on mESC (11). **(B)** *Cis* binning strategy with geometrically increasing bins provides more even contact coverage in the vicinity of the source gene.

Finally, functionally diverse classes of RNAs have different patterns of genome occupancy. For instance, an enhancer RNA – promoter interaction may exhibit a narrow peak profile, while a small nuclear RNA can appear to be broadly distributed along gene bodies. Furthermore, we expect that peak width would be affected by the amount of the data, which refers to resolution.

In ATA datasets, RNAs differ in contact frequency, while limited sequencing depth leads to data sparsity as a result of incomplete contact capture. Consequently, there is a ground to introduce a variable peak size for different RNAs depending on their abundances and distribution patterns along the chromatin. Existing GRID and RADICL procedures propose fixed-size bins for all RNAs, which can reduce the resolution for RNAs with narrow binding sites, and lower the statistical power for RNAs with more diffuse binding sites.

In this work, we introduce a versatile tool BaRDIC (Binomial RNA-DNA Interaction Caller), that utilizes a binomial model to identify genomic regions significantly enriched in RNA-chromatin interactions, or “peaks”, in ATA and OTA data. Our approach accounts for several key features of RNA-DNA interaction data:

- chromatin heterogeneity;
- scaling;
- data sparsity and distinct RNA contact patterns.

BaRDIC combines approaches from Hi-C and ChIP-seq data analysis and yields accurate predictions of RNA-DNA interactions from both OTA and ATA data.

## MATERIALS AND METHODS

### Peak-calling on OTA data

Pre-processed contacts and MACS2 peaks for OTA experiments were extracted from the RNA-Chrom database (17). The data identifiers (Exp. ID) are as follows: 94 (RAP-RNA Malat1), 102 (ChIRP Halr1) for mESC cell line, 90 (CHART Paupar) for mouse neuroblastoma cell line N2A. Details on the experiments and data processing protocol are described in the RNA-Chrom database.

Background track for OTA experiments was constructed from input data and converted to begGraph format in 1kb bins using bedtools package (20). The type of background used by BaRDIC was set to a pre-constructed background track (-bt custom). Due to relatively high resolution of OTA experiments, the following parameters were provided for peak-calling on OTA data: 400 nt minimum *trans* bin size (– trans min), 100 nt initial *cis* bin size (–cis start 100), 50 nt step to find the optimal *trans* bin size (–trans step). For Halr1 (Haunt), the peaks for the retinoic acid-induced mESC sample were used for analysis as the peaks from the control mESC sample do not intersect HOXA cluster genes.

### Peak-calling on ATA data

Contacts for the GRID-seq ATA experiment on the mESC cell line (Exp. ID 6) and RNA gene annotation for the mouse genome (except X-RNA biotype) were extracted from the RNA-Chrom database. Peaks were called with default parameters. Peaks by RADICL procedure were called according to the algorithm description from the original study (16). Since we did not succeed in reproducing the original GRID peak-calling method, the peaks published by authors were used in analysis (11). To compare BaRDIC with the GRID peak caller, the publicly available peaks were converted from mm9 to mm10 using LiftOver (21). It is worth noting that the peak comparison may be skewed due to differences in the data processing protocol of the GRID-seq authors and our group.

## RESULTS

### BaRDIC algorithm

The BaRDIC algorithm consists of three steps:

1. Binning: for each RNA, chromosomes are partitioned into non-overlapping genomic intervals (bins), and the number of contacts is calculated within bins.

2. Statistical modeling: for each RNA, parameters of the background model are calculated in every bin. P-values are calculated based on them.

3. Multiple testing correction with Benjamini-Hochberg procedure (22).

To increase the statistical power and speed up the algorithm’s performance, only RNAs with more than 1000 contacts are selected by default. Steps 1 and 2 are atomic with regard to individual RNAs, so we describe them for one RNA only.

*Binning*. Bin sizes are estimated for each RNA separately. The binning strategy differs for *cis* interactions, which occur on a chromosome harboring the RNA gene, and *trans* interactions – with other chromosomes.

In the case of random ligation, we expect that *trans* contacts are distributed uniformly along chromosomes. Therefore, we apply a uniform binning strategy for *trans* interactions.

For binning in *cis*, scaling must be taken into account. Also, we assume long-range *cis* interactions are similar to *trans* interactions, analogous to observations in Hi-C data analysis (23). In our *cis* binning strategy, chromosomes are partitioned into non-uniform bins of size increasing in a geometric progression from the source gene. *Cis* bin size increases until it exceeds the *trans* bin size; all subsequent *cis* bins are uniform and equal to the *trans* bin in size (see Figure 1B):

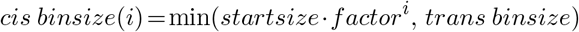

To find optimal bin sizes, we adapted RSEG (24) and JAMM(25) approaches that were developed for ChIP-seq peak-calling. This approach finds a balance between resolution and sufficient bin coverage.

*Statistical modelling*. To model the background distribution of RNA-DNA contacts in bins, we introduce a frequentist model similar to those used in ChIP-seq and Hi-C data analysis (26, 27). Assuming that contacts arising from random binding are independent, we consider the number of contacts *X*_*ij*_ of RNA *i* in bin *j* to be binomially distributed:

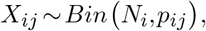

where *N*_*i*_ is the total number of contacts produced by RNA *I* (except the gene body), *p*_*ij*_ is the background probability.

Note the number of observed contacts *O*_*ij*_ is a realization of the random variable *X*_*ij*_. Statistical estimation comes down to inferring the only model parameter *p*_*ij*_ from the observed data.

For *trans* bins, only chromatin heterogeneity plays a role. In ATA experiments, we estimate the parameter of the background model by counting mRNA *trans* contacts as proposed in the GRID procedure. The background probability of a single contact of RNA *i* to appear in the *j*-th *trans* bin is assumed to be as follows:

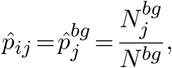

Where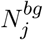 is the number of contacts from the background in a bin *j, N*_*bg*_ is the total number of background contacts. For OTA experiments, we use contacts from the input sample similar to ChIP-seq analysis.

For *cis* bins, we additionally consider scaling. To do that, we define a scaling factor *f* (*d*_*ij*_) that depends on the distance between the source gene of RNA *i* and bin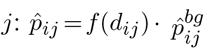 We estimate the scaling factor as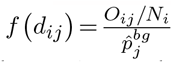 and smooth its profile using a cubic smoothing spline in double logarithmic coordinates. As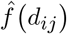 values vary substantially for different RNAs, splines are calculated for every RNA separately. To exclude putative *cis* peaks from the background model, we perform a two-step scaling estimation procedure similar to Fit-Hi-C (26) and HiC-DC (28), removing bins with statistically high contact coverage.

To hold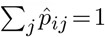, we renormalize probability estimates for each RNA *i*. As a result, the sum of background probabilities for *cis* bins equals the fraction of *cis* contacts of each RNA, the same is true for *trans* bins.

We expect that a specific binding event results in a high enrichment of contacts relative to the background. The resulting P-value is then computed with the right-sided binomial test using estimated background parameters. P-values from non-zero bins are subjected to the multiple testing correction using the Benjamini-Hochberg procedure, simultaneously for all RNAs.

*Implementation*. The algorithm is implemented as a package for Python 3 (29) with a command line interface and is available at https://github.com/dmitrymyl/BaRDIC. It is based on numpy (30), pandas, scipy (31) and statsmodels packages (32). The bioframe package (33) is used for operations with genomic intervals. Since binning and statistical evaluation are performed for each RNA separately, these steps are parallelized.

To organize the data storage and speed up the data access, we developed two HDF5-based data formats (34), which were inspired by the cool (35) format. Detailed algorithm description and format schemas are given in Supplementary Note.

### Overview of evaluation and use cases

Evaluating the peak caller performance on RNA-DNA interaction data is still an open problem, given the lack of a gold standard or a “true” dataset. To address this, we will assess peak calling results based on biological principles of RNA-chromatin interactions:

1. We assess the quality of BaRDIC peaks based on well-studied RNAs with known binding preferences;
2. We compare BaRDIC performance with the three current approaches to identify RNA-binding sites — MACS2 peak caller for OTA and original methods from ATA studies, GRID-seq and RADICL-seq, hereafter referred to as GRID-peak and RADICL-peak.

Importantly, these three current approaches do not model all typical biases in RNA-DNA interaction data, e.g. scaling, so none of them can be considered the gold standard.

### General assessment of BaRDIC ATA peaks

We evaluated BaRDIC by using GRID-seq data on the mESC, a cell line for which many OTA experiments are also available (See Supplementary Tables S1, S2). Only RNAs with at least 1000 contacts in the ATA experiment were selected for peak calling. In total, BaRDIC identified specific interactions mediated by 7741 unique RNAs in GRID-seq data on mESC (Supplementary Figure S1, S3). For the majority of RNAs, the size of *trans* bins ranges from 30 to 80 Kb, with a median of 50 Kb (Supplementary Figure S2). RNAs of distinct biotypes were found to have different preferences for *cis* and *trans* interactions. In general, small non-coding RNAs have a large fraction of *trans* peaks, while long non-coding RNAs can generate both *cis* and *trans* peaks (Figure 2A). Notably, RNA preferences do not correlate with the number of RNA peaks, which suggests BaRDIC is robust against data sparsity.

**Figure 2.**
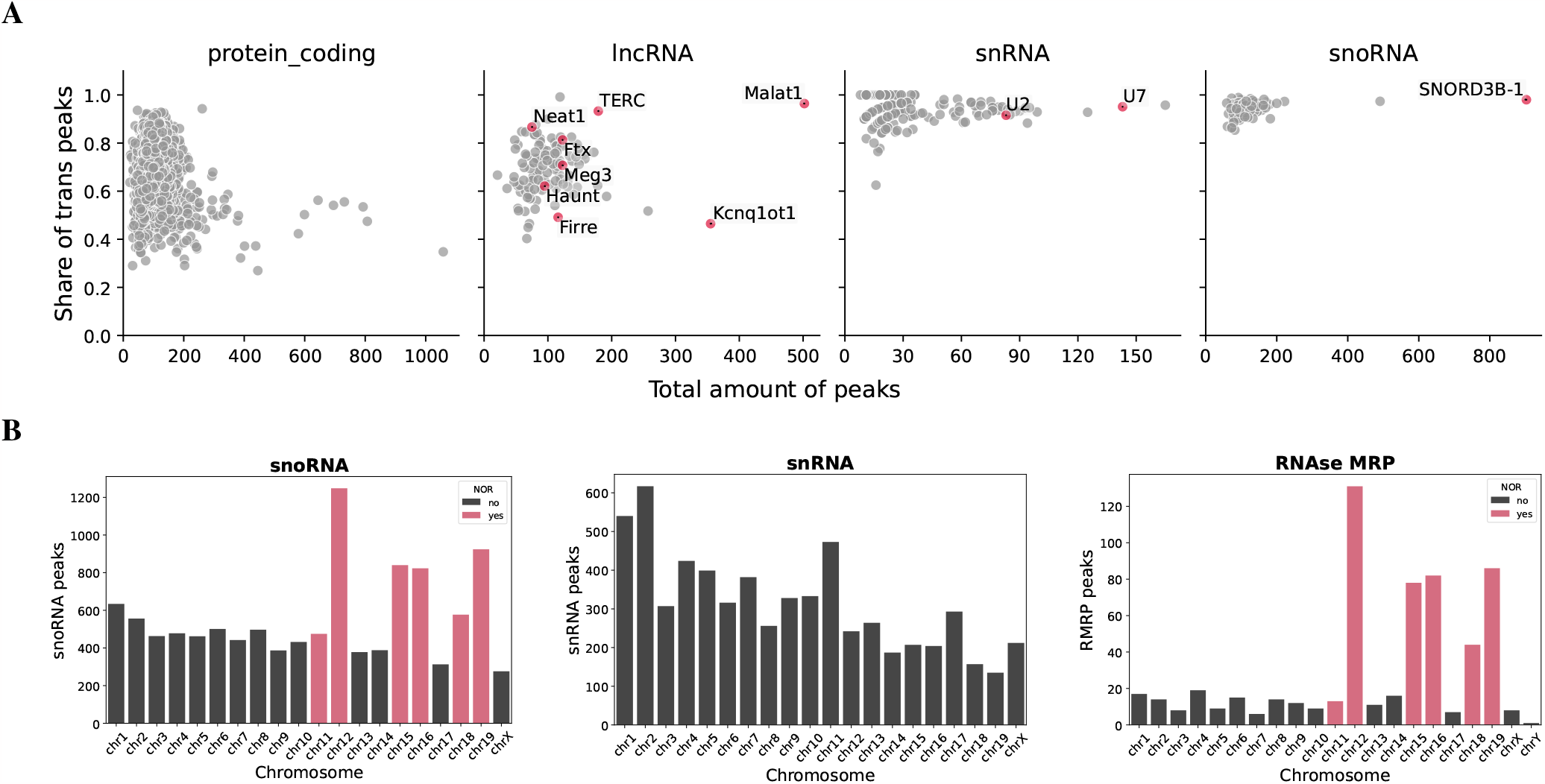
Global assessment of specific RNA interactions identified by BaRDIC for GRID data on mESC data. **(A)** Fractions of *trans* peaks and the number of ATA peaks for select RNA biotypes. **(B)** Distribution of snoRNAs, snRNAs, and MRP RNase peaks across mouse chromosomes. Chromosomes carrying nucleolus organizer regions (NORs) are highlighted in blue.

For further analysis, 1007 unique RNAs were selected: all RNAs with at least one BaRDIC peak, except for ribosomal RNAs and mRNAs. To generally assess specific interactions in ATA data, we focused on RNAs with well-studied binding patterns. As expected, Malat1, SNORD3B-1, TERC, Neat1, and RMRP prefer to contact in trans, while Kcnq1ot1, Firre, and Haunt (Halr1) favor *cis* interactions (Figure 2A, Supplementary Figure S4). Long ncRNAs as a class have variable preferences because they can act both as *cis* and *trans* regulators (36). SnoRNAs generate many *trans* peaks as they interact with chromosomes carrying the nucleolus organizer regions (NORs): chromosomes 12, 15, 16, 18, 19 except for chromosome 11 (Figure 2B) (37). A similar distribution is observed for RNase MRP (RMRP) consistent with its localization in the nucleus (38). Apparently, in this particular mESC sample, chromosome 11 does not carry active NOR loci. In contrast, snRNAs do not have specific chromosomal preferences: the number of peaks on chromosomes correlates with their lengths (*r*_*S*_ = 0.71, *p− value<* 3.1*e−* 4).

Taken together, the coincidence of *cis* and *trans* peaks distribution with known RNA binding preferences validates BaRDIC peaks at the coarse-grain level.

### Detailed comparison of peaks of individual RNAs

To assess the quality of BaRDIC peaks at the local level, we obtained peaks for several RNAs with known binding patterns, for which OTA experiments are available (Supplementary Tables S1, S2). When comparing OTA and ATA experiments for a particular RNA, it is crucial to keep in mind that ATA contacts are much sparser (Supplementary Figure S5) and located at varying distances from defined OTA peaks due to the differences in experimental procedures (Supplementary Figure S6). Therefore, in addition to the direct comparison of peaks, we compare genes intersecting peaks, similar to the RADICL approach (16).

To highlight features of BaRDIC, we chose three non-coding RNAs (ncRNAs) with distinct distributions of contacts. ncRNA Paupar exhibits virtually no scaling; ncRNA Malat1 is abundant in ATA datasets and exhibits a pronounced scaling; ncRNA Halr1 is unabundant RNA in ATA datasets, regulates gene expression in *cis* and therefore should have peaks very close to the source gene.

### Paupar: ncRNA with no scaling

Paupar is a vertebrate-conserved lncRNA whose expression is restricted to the central nervous system. CHART-seq revealed that Paupar is predominantly associated with promoters and 5’-UTR regions of protein-coding genes and is almost evenly distributed across chromosomes, except for the lack of contacts on the X chromosome. We have shown that the scaling effect is not pronounced for Paupar (Supplementary Figure S7E). Thus, we can directly compare the performance of BaRDIC and other algorithms given the virtually absent influence of scaling. As ATA data for mESC lacks Paupar interactions, we compared only OTA peaks.

Sets of genes that intersect BaRDIC peaks and MACS2 peaks for OTA were very similar (Figure 3A). We confirmed spatial co-localization of BaRDIC and MACS2 peaks using GenometriCorr (39) metrics (Figure 3B; Supplementary Figure S8). Therefore, for an RNA with a diffuse genome-wide distribution and minor scaling effects, BaRDIC peaks are in agreement with the publicly available results. This proves BaRDIC captures chromatin heterogeneity similarly to other peak calling algorithms.

**Figure 3.**
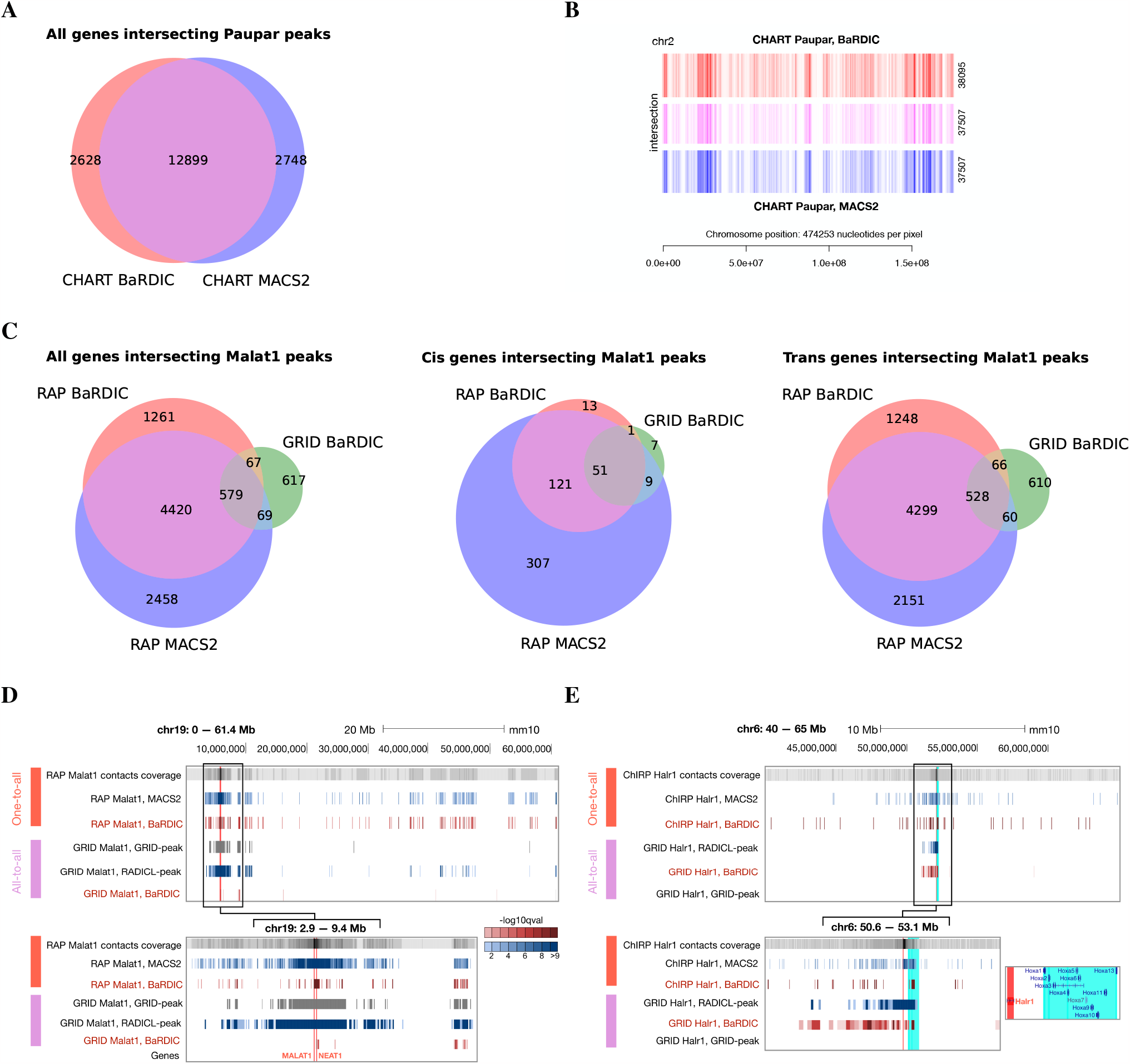
Comparison of peaks obtained by different algorithms for OTA and ATA data on mESC cell line **(A)** Genes intersecting Paupar peaks from CHART data obtained by BaRDIC and MACS2. **(B)** Global patterns of peak distribution and peaks intersections on “parental” chromosome 2, CHART data. **(C)** Genes intersecting Malat1 peaks from RAP and GRID mESC data. From left to right: all genes, *cis* genes, *trans* genes. **(D)** Representative genome browser view of Malat1 peaks identified by different algorithms in RAP OTA data (red) and ATA data (purple). Scales are given: the entire length of mouse chromosome 19 and the vicinity of the Malat1 gene on chromosome 19. Vertical red lines represent Malat1 and Neat1 source genes. **(E)** Representative genome browser view of Halr1 peaks identified by different algorithms in ChIRP OTA data (red) and ATA data (purple). The Halr1 gene located on chromosome 6 is highlighted in red, HOXA gene cluster in its vicinity is highlighted in cyan. Tracks are colored by *−log*_10_*Q* statistical significance when appropriate.

### Malat1: abundant ncRNA with pronounced scaling

Malat1, an abundant lncRNA, is a global regulator of transcription and pre-mRNA splicing. Malat1 has multiple binding sites in *trans*, predominantly in the bodies of actively transcribed genes (40). Moreover, its possible *cis* regulatory role (41, 42) in controlling Neat1 expression has been discussed. Remarkably, Malat1 generates over 527,000 contacts in the selected ATA experiment, being the RNA with the largest number of contacts (Supplementary Table S1; Supplementary Figure S9). Given a sufficient number of Malat1 *cis* contacts, we investigated the influence of the scaling effect (Supplementary Figure S7A, B) on the performance of peak calling algorithms.

BaRDIC on the OTA data reproduces the set of genes that intersect MACS2 peaks (Figure 3C), in addition, peaks are spatially co-localized (Supplementary Figure S10). Importantly, BaRDIC peaks are more evenly distributed along chromosome 19 which harbors the Malat1 source gene. MACS2 does not take into account scaling and overestimates the significance of peaks near the Malat1 gene boundaries, which is corrected by the BaRDIC algorithm (Figure 3D).

When compared with GRID-peak, BaRDIC shows partial reproducibility of genes intersecting peaks (Supplementary Figure S11). Genes intersecting Malat1 *cis* peaks called with BaRDIC form a subset of those intersecting peaks called with GRID-peak. GRID-peak and RADICL-peak calls are clustered near the Malat1 source gene (Figure 3D). This suggests that BaRDIC filters out *cis* peaks that arose from scaling but preserves previously described specific interactions at the Neat1 locus (40).

Due to scaling correction, BaRDIC improves previous strategies as it does not overestimate the statistical significance of sites proximal to the RNA source gene. At the same time, BaRDIC preserves consensus target genes, which are likely to be true targets of chromatin-associated RNAs.

### Halr1: cis-acting unabundant ncRNA

Long ncRNA Halr1 (HOXA upstream noncoding transcript) regulates the HOXA gene cluster during ESC differentiation (43). In the mouse genome, the HOXA gene cluster is located _*∼*_40 Kb downstream of the Halr1 transcription site. Halr1 is an unabundant RNA in ATA experiments, with 1724 contacts (23rd percentile) (Supplementary Table S1; Supplementary Figure S9) and pronounced scaling (Supplementary Figure S7C, D) in GRID mESC data. We therefore included a Halr1 case study to examine the quality of the BaRDIC algorithm on low-contacting RNAs, which form the vast majority of ATA data.

Peaks called with BaRDIC in OTA and ATA data indeed covered the genes of the HOXA gene cluster (Figure 3E). In contrast, GRID-peak identifies no Halr1 peaks. Additionally, the geometric binning procedure in *cis*, implemented in BaRDIC, increases the resolution near the boundaries of the Halr1 gene (Supplementary Figure S12). This results in a more accurate localization of peaks near the RNA source gene.

Overall, BaRDIC outperforms other peak callers in detecting peaks of RNAs with a small number of contacts in the ATA data. In addition, despite accounting for scaling, BaRDIC finds biologically relevant specific interactions near the RNA source gene.

## DISCUSSION

In this paper, we present BaRDIC, a first-of-its-kind versatile algorithm for identifying specific RNA-chromatin interactions in both “one-to-all” (OTA) and “all-to-all” (ATA) data. BaRDIC takes into account typical biases and other features of RNA-DNA interaction data, including chromatin heterogeneity, scaling, and distinct contact patterns of individual RNAs. In the virtual absence of scaling effects, BaRDIC performs comparably with other algorithms. In the presence of scaling, BaRDIC does not overestimate the statistical significance of findings near the source gene of an individual RNA, unlike previously published approaches. The adaptive bin size selection for each RNA and geometric binning near source genes in ATA data allows for more accurate identification of specific binding sites. In addition, BaRDIC is robust to data sparsity, which is a hallmark of ATA experiments.

We found that when calling peaks in OTA data, BaRDIC and the commonly used MACS2 peak caller incompletely reproduce each other’s results on specific *trans* interactions, even with no apparent influence of scaling. This may be due to different strategies of genome binning: BaRDIC relies on non-overlapping fixed-size bins, while MACS2 determines regions (peak candidates) dynamically. Perhaps the combined approach of binning with offsets in BaRDIC would increase the reproducibility of the results. In addition, the discrepancies may arise from different statistical models — a dynamic Poisson model in MACS2 and a binomial model in BaRDIC. Finally, BaRDIC peaks for RNAs in this study are wider than MACS2 peaks (Supplementary Table S2). Consequently, BaRDIC bins may include more non-specific interactions, which also decreases the resolution. Therefore, we controlled FDR more stringently for BaRDIC findings compared to MACS2. However, it is not our goal to achieve the full reproducibility of results between BaRDIC and MACS2 in OTA data, as there is currently no gold standard for specific RNA interactions. Besides, MACS2 does not account for scaling – the prominent bias in RNA-DNA interaction data, particularly in OTA data. Nevertheless, partial, but substantial reproducibility of peaks implies that BaRDIC accounts for chromatin heterogeneity similarly to other algorithms used in this area of research.

For all RNAs studied in this paper, peaks from ATA experiments intersect only a fraction of genes corresponding to OTA experiments, and may as well capture some non-target genes. This may be due to the insufficient sequencing depth in ATA experiments and, as a consequence, dropout. In this process, many true RNA binding sites are lost before the sequencing step, and the remaining nonspecific sites distort the statistical significance of detected peaks. This assumption is supported by the fact that the reproducibility of target genes is better for RNAs with a higher number of contacts in ATA data, such as Malat1. Differences in experimental procedures may also contribute to the inconsistency between OTA and ATA peaks (Supplementary Figure S6). OTA experiments are similar to ChIP-seq with a DNA shearing step that allows a relatively accurate identification of a contact region. In contrast, ATA experiments are similar to Hi-C experiments with DNA restriction and proximity ligation steps, which limit observed contact regions to restriction sites. Collectively, OTA and ATA peaks fundamentally cannot be compared properly, so we do not expect an exact match of called interactions due to the differences and limitations of the two types of experiments.

In this work, we highlighted the importance of the scaling effect when detecting RNA peaks on chromatin, which has already been around in the field. A recent preprint on ChAR-seq (44) proposes a generative model for predicting peak-like interactions based on mRNA *trans* contacts, with an account for scaling. However, that model suggests fix-sized bins for all RNAs, while BaRDIC proposes bins of variable size, individually for each RNA. This feature allows BaRDIC to model non-uniform contact coverage and take into account distinct distribution patterns of RNAs along the chromatin.

BaRDIC is not only applicable to pairwise RNA-DNA interaction data but can also be applied to multiway RD-SPRITE (45) data. To achieve this, the multiway data needs to be expanded into virtual pairwise RNA-DNA contacts similar to the strategy proposed for analyzing multiway DNA-DNA interactions obtained with Pore-C (46).

BaRDIC package is an easy-to-use tool since it requires only three files as input: a BED file with DNA parts of contacts, an RNA gene annotation, and information on the background — a list of background mRNAs or an input track. This simplicity allows a user to parallelize peak calling over multiple RNAs with an ATA dataset or streamline BaRDIC application over multiple isolated datasets. Calling peaks on all available OTA and ATA experiments with scaling correction could help to refine known target genomic loci for each RNA of interest. Collectively, the resulting peaks can be organized in a database, such as RNA-Chrom. The unified approach will enable unbiased comparisons of specific RNA-DNA interactions between samples and studies as well as with other high throughput omics data.

Scaling correction during peak calling makes RNA-DNA interaction data reconcilable with specific protein-DNA interactions (ChIP-seq peaks), for which scaling does not exist, and specific DNA-DNA interactions (Hi-C loops), for which scaling is observed and accounted for in dedicated algorithms. This harmonization opens a way for unbiased comparisons of these three types of interactomic data for predicting functions of RNA-chromatin interactions. In particular, these comparisons will lead to predictions about how RNAs bind to their genomic targets: either due to a spatial proximity of corresponding DNA loci or due to recruitment by DNA-binding proteins. In turn, GO and GSEA enrichment analysis of genes interacting with novel chromatin-associated RNAs will create a basis for predicting regulatory functions of RNA-chromatin interactions. Taken together, simultaneously accounting for chromatin heterogeneity and scaling in RNA-DNA interaction data enables unbiased and uncomplicated downstream and comparative analyses.

## ACKNOWLEDGEMENTS

We would like to thank G. Ryabykh for valuable suggestions and feedback on the manuscript. The research was carried out using facilities of the Makarich HPC cluster provided by the Faculty of Bioengineering and Bioinformatics, Lomonosov Moscow State University.

