## Supplementary Data for "BaRDIC: robust peak calling for RNA-DNA interaction data"

| RNA | Cell line | Total number of contacts | BaRDIC *trans* bin size, Kb | Amount of BaRDIC peaks | Amount of GRID peaks | Amount of RADICL peaks |
|---|---|---|---|---|---|---|
| Malat1 | mESC | 527 249 | 20 | 502 | 5042 | 2515 |
| Halr1 | mESC | 1724 | 43 | 96 | 0 | 24 |

**Supplementary Table S1:** Number of contacts in ATA GRID data in mESC cell line and the amount of peaks called with different methods — BaRDIC, GRID-peak, and RADICL-peak. Selected *trans* bin size for BaRDIC is shown; GRID-peak and RADICL-peak bin sizes are fixed and equal 1 Kb and 25 Kb, respectively.

| RNA | OTA experiment | Cell line | Total number of contacts | BaRDIC *trans* bin size, bp | Amount of BaRDIC peaks | Amount of MACS2 peaks | Median peak size, MACS2 |
|---|---|---|---|---|---|---|---|
| Malat1 | RAP | mESC | 14 417 923 | 900 | 14 166 | 15 962 | 394 |
| Paupar | CHART | N2A | 59 369 258 | 400 | 78 996 | 68 833 | 295 |
| Halr1 | ChIRP | mESC | 4 772 537 | 5350* | 2 835 | 2 069 | 178 |

**Supplementary Table S2:** Number of contacts and amount of peaks called with BaRDIC and MACS2 for OTA data. As OTA peaks called with BaRDIC are generally wider than those called with MACS2, we applied more stringent thresholds to select more prominent BaRDIC peaks: $-log10$(q-value)$>$ 4 for Malat1, $-log10$(q-value) $>$ 7 for Paupar and Halr1. *Initial *cis* bin size equals 100, Halr1 preferentially interacts in *cis*.

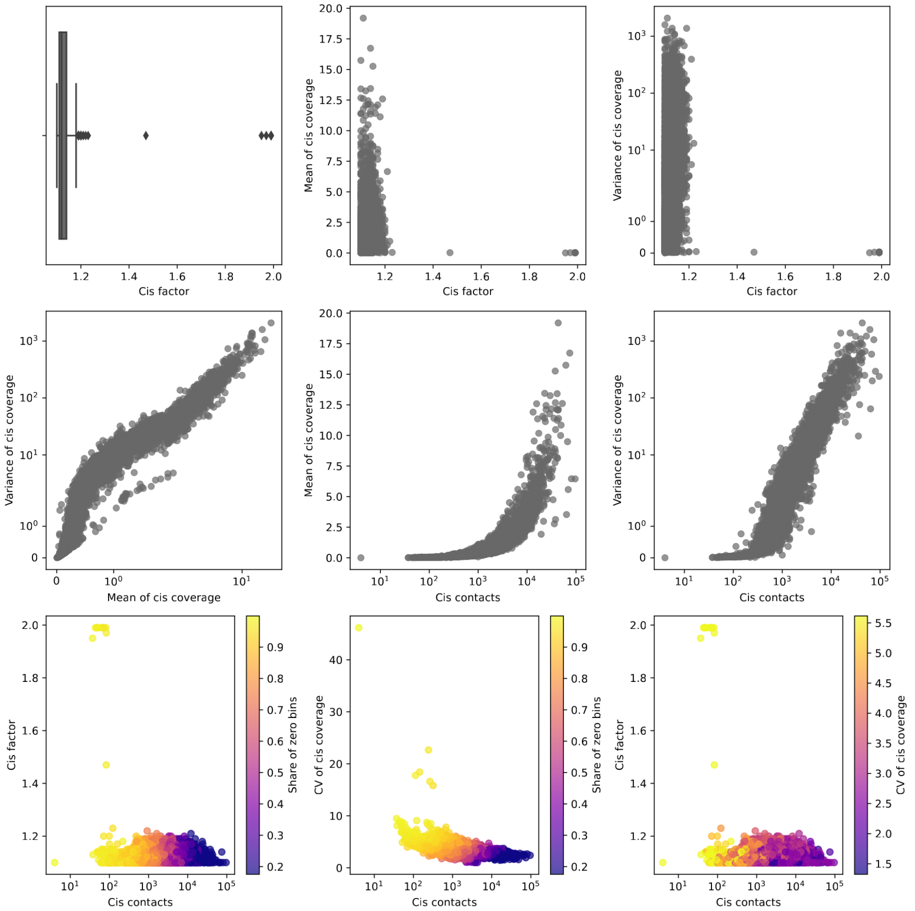


**Supplementary Figure S1:** Results of *cis factor* selection for RNAs from GRID data on mESC cell line. Only RNAs with more than 1000 contacts were considered. From left to right, from top to bottom: (1) *cis* factor distribution; the dependence of average *cis* bin coverage (2) and contact coverage variance (3) on bin size; 4) the ratio of average *cis* bin coverage by contacts to variance; the dependence of average *cis* bin coverage (5) and coverage variance (6) on the number of *cis* contacts; 7) the dependence of *cis* bins size on the number of *cis* contacts with a fraction of zero bins encoded by color; 8) dependence of the coefficient of variation (CV) of *cis* bins coverage on the number of *cis* contacts, a fraction of zero bins encoded by color; 9) the dependence of *cis* bin size on the number of *cis* contacts, CV of coverage encoded by color.

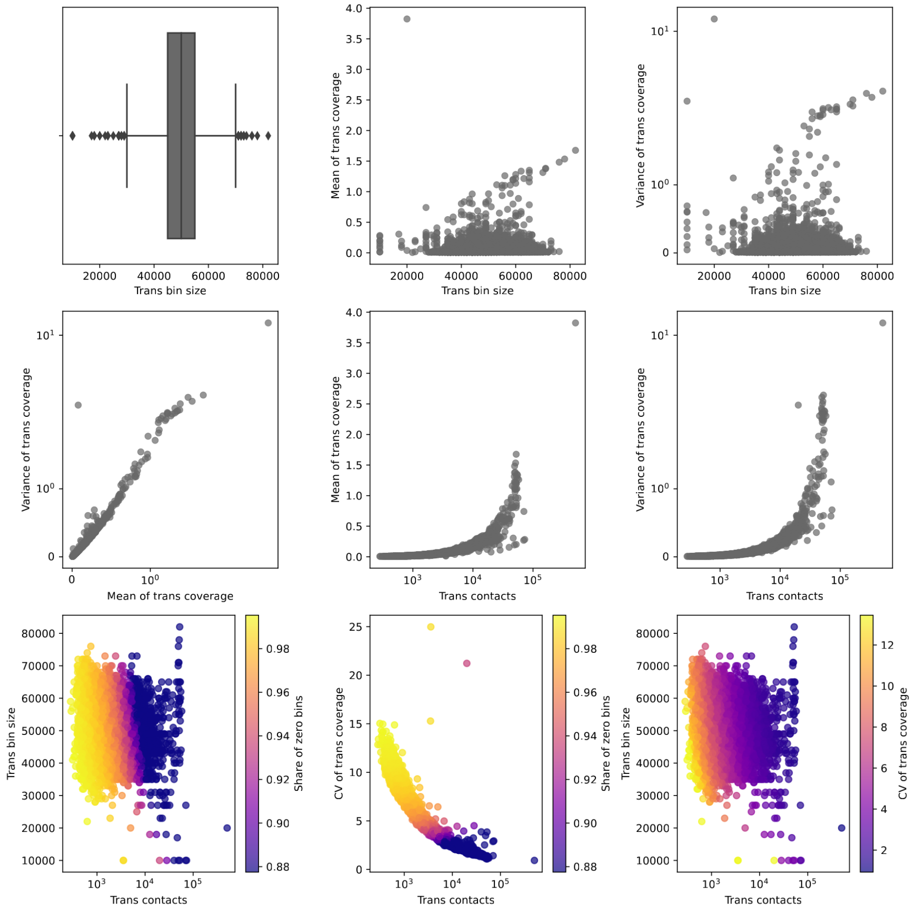


**Supplementary Figure S2:** Results of *trans bin* size selection for RNAs from GRID data on mESC. Only RNAs with more than 1000 contacts were considered. From left to right, from top to bottom: (1) *trans* bin size distribution; the dependence of average *trans* bin coverage (2) and contact coverage variance (3) on bin size; 4) the ratio of average *trans* bin coverage by contacts to variance; the dependence of average *trans* bin coverage (5) and coverage variance (6) on the number of *trans* contacts; 7) the dependence of *trans* bins size on the number of *trans* contacts with a fraction of zero bins encoded by color; 8) dependence of the coefficient of variation (CV) of *trans* bins coverage on the number of *trans* contacts, a fraction of zero bins encoded by color; 9) the dependence of *trans* bin size on the number of *trans* contacts, CV of coverage encoded by color.

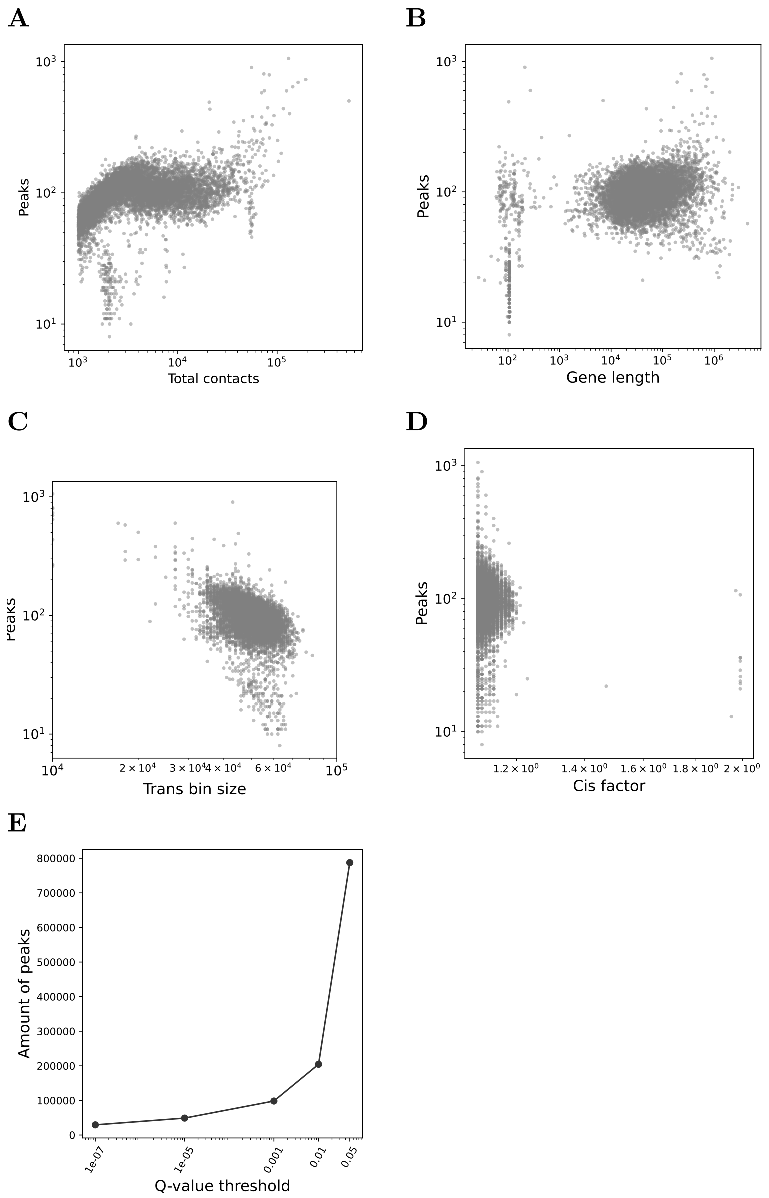


**Supplementary Figure S3:** Exploration of peaks called with BaRDIC from GRID data on mESC. Dependency of the number of peaks on (A) the total number of RNA contacts ($r_S = 0.45$), (B) the length of RNA source gene ($r_S = 0.23$), (C) selected *trans* bin size ($r_S = -0.44$), and (D) *cis* factor ($r_S = -0.02$) are shown. (E) Numbers of peaks at different Q-value thresholds, including mRNAs peaks.

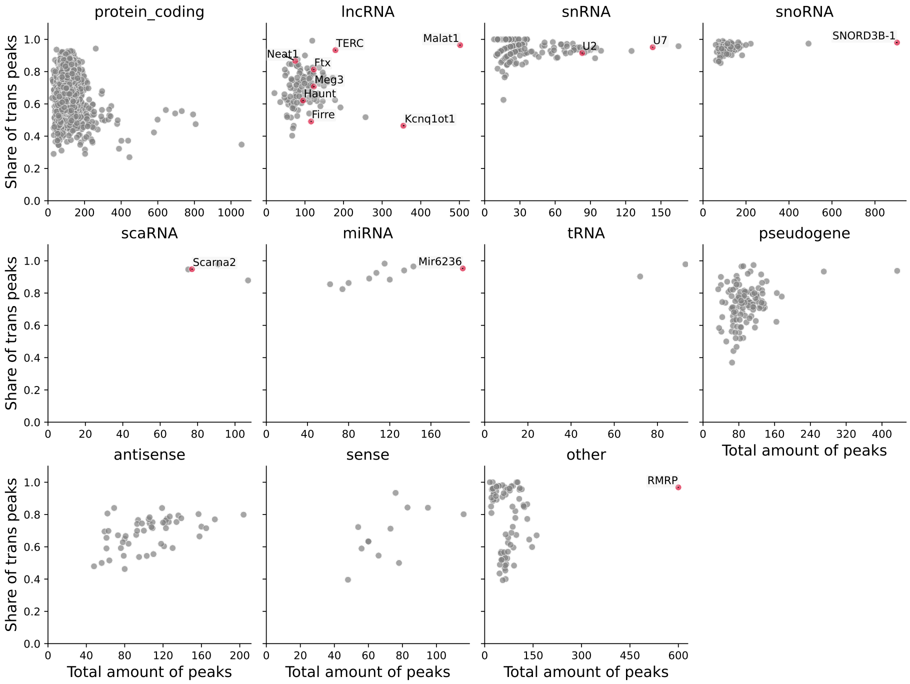


**Supplementary Figure S4:** Global assessment of specific RNA interactions identified by BaRDIC for GRID data on mESC data. Fractions of *trans* peaks and the number of ATA peaks for different RNA biotypes.

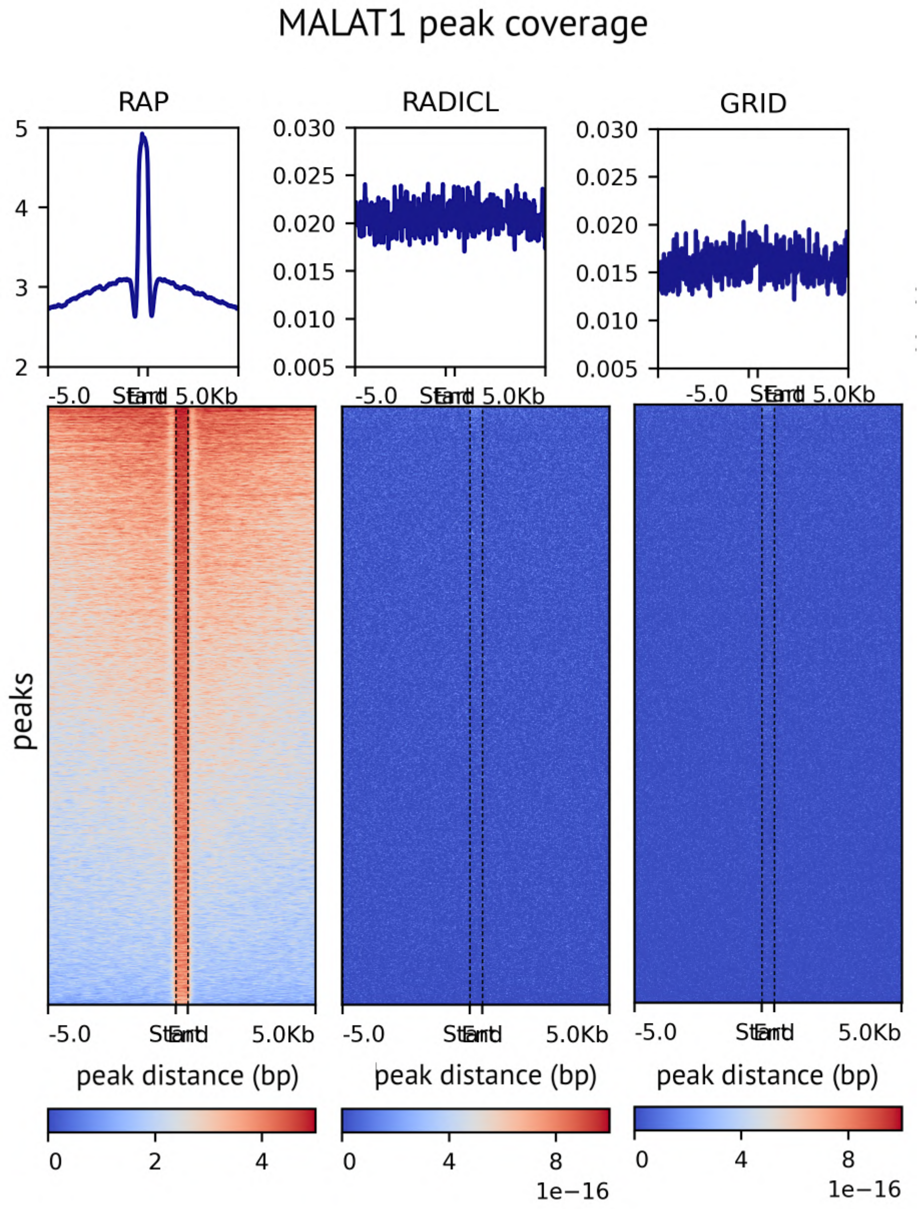


**Supplementary Figure S5:** MACS2 peaks for Malat1 RAP OTA data and contacts from RAP, RADICL and GRID for mESC. The coverage of OTA peaks with ATA contacts for mESC.

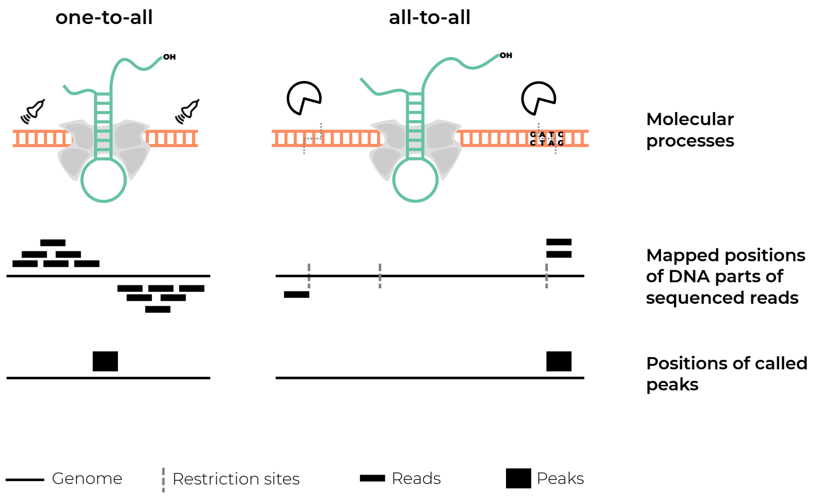


**Supplementary Figure S6:** For OTA experiments with DNA shearing, read form groups on both strands around the binding site. For all-to-all experiments with DNA digestion, DNA parts of contacts map to restriction sites downstream in 5' direction.

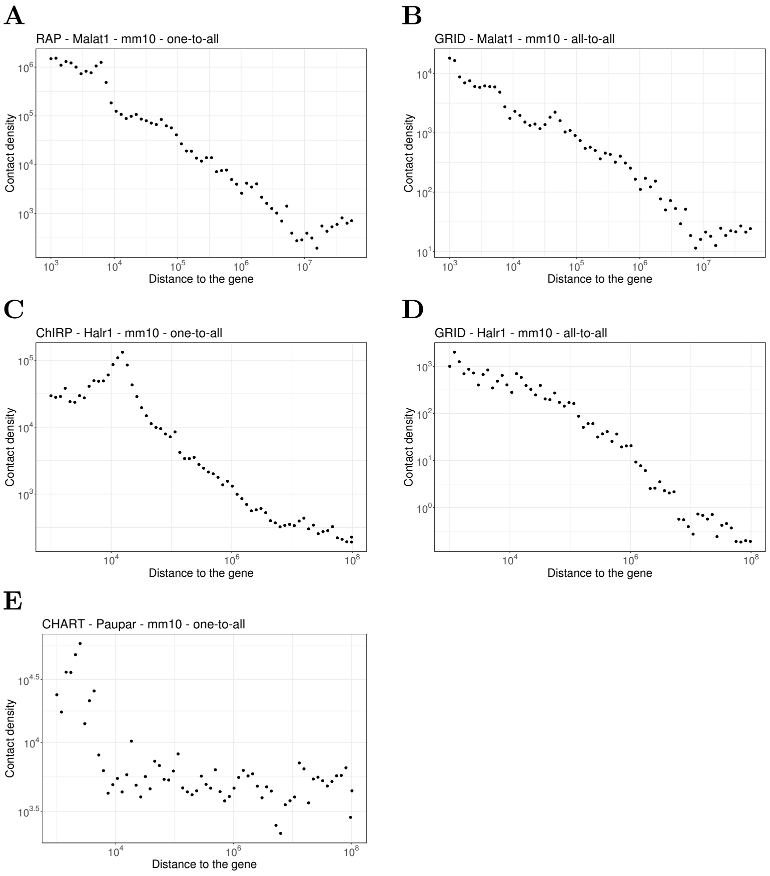


**Supplementary Figure S7:** Dependency of contact density on the distance between the RNA source gene and chromatin target loci (scaling) in double logarithmic coordinates. **(A)** Scaling of Malat1 in OTA RAP data, mESC **(B)** Scaling of Malat1 in ATA GRID data, mESC **(C)** Scaling of Halr1 in OTA ChIRP data, mESC. **(D)** Scaling of Halr1 in ATA GRID data, mESC. **(E)** Scaling of Paupar in OTA CHART data, N2A cell line.

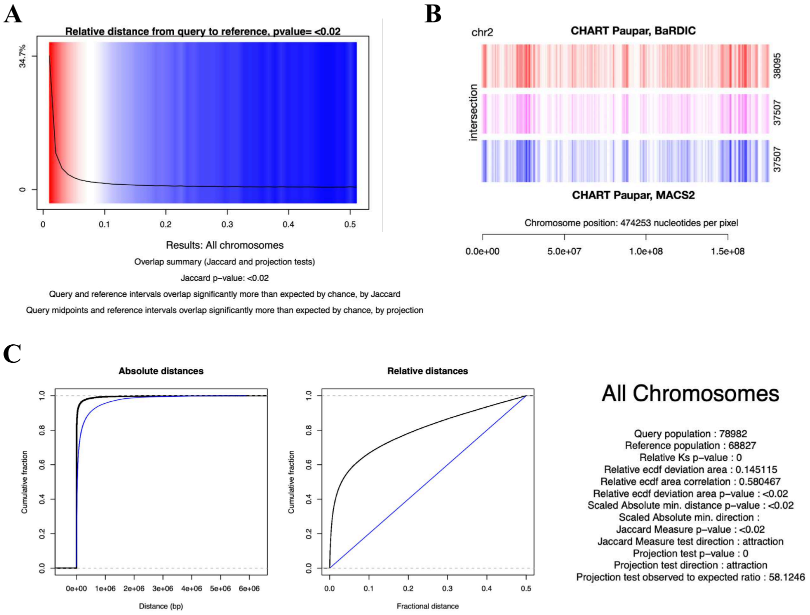


**Supplementary Figure S8:** GenometriCorr was used to examine pairwise spatial correlations between genomic profiles: BaRDIC peaks (Query) and MACS2 peaks (Reference) called from CHART Paupar data. GenometriCorr implements various statistical approaches that are based on interval overlaps, relative genomic distances, or absolute genomic distances. In absolute and relative distance tests, intervals are represented by their midpoints. (A) Areas of high and low relative distance correlation. Red and blue colors indicate deviation from the expected distribution, while the black line indicates the density of the data in these regions. (B) Graphical representation of BaRDIC and MACS2 peaks for chromosome 2 harboring source gene. (C) Statistical summary and ECDF plot for the relative and absolute distances for the entire genome. The blue line represents the expected distribution (no association), and the black line represents the actual distribution of data. Relative distance tests show a correlation, this indicates that BaRDIC and MACS2 peaks called for Paupar tend to co-localize.

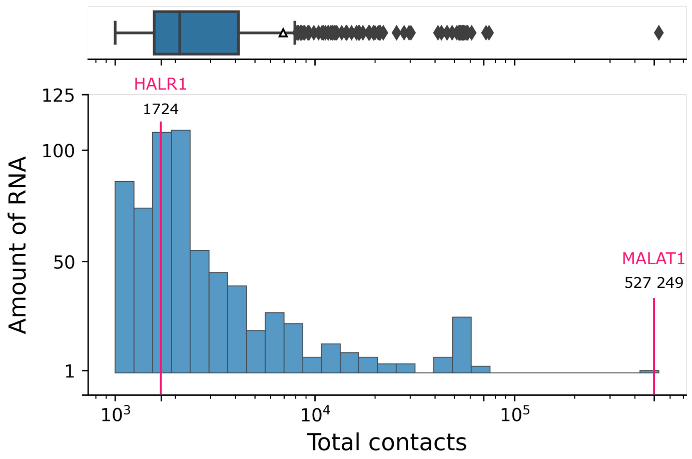


**Supplementary Figure S9:** The distribution of number of contacts for all RNAs ($> 1000$ contacts) in GRID data on mESC. Ribosomal RNAs and mRNAs are excluded. Individual RNAs are shown: Malat1 as an example of an abundant RNA, Halr1 as an example of an unabundant RNA.

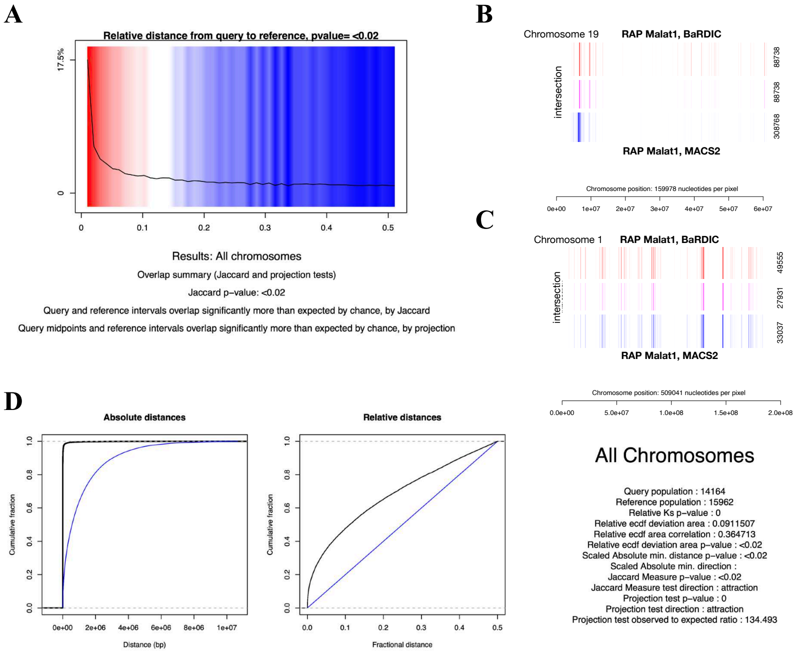


**Supplementary Figure S10:** GenometriCorr was used to examine pair-wise spatial correlations between genomic profiles: BaRDIC peaks (Query) and MACS2 peaks (Reference) called from RAP Malat1 data. (A) Areas of high and low relative distance correlation. Red and blue colors indicate deviation from the expected distribution, while the black line indicates the density of the data in these regions. (B) Graphical representation of genomic profiles for chromosome 19 harboring Malat1 source gene. (C) Graphical representation of BaRDIC and MACS2 peaks for "non-parental" chromosome 1 (D) Statistical summary and ECDF plot for the relative and absolute distances for the entire genome. The blue line represents the expected distribution (no association), and the black line represents the actual distribution of data. Relative distance tests show a correlation, this indicates that BaRDIC and MACS2 peaks for Malat1 tend to co-localize.

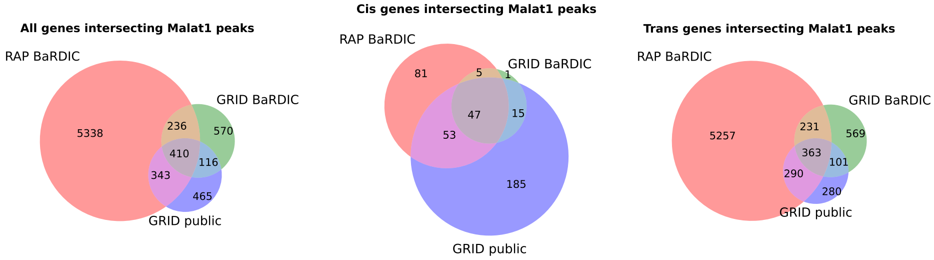


**Supplementary Figure S11:** Genes intersecting Malat1 peaks from RAP and GRID mESC data. BaRDIC OTA and ATA peaks and GRID-peak ATA peaks are compared. From left to right: all genes, *cis* genes, *trans* genes.

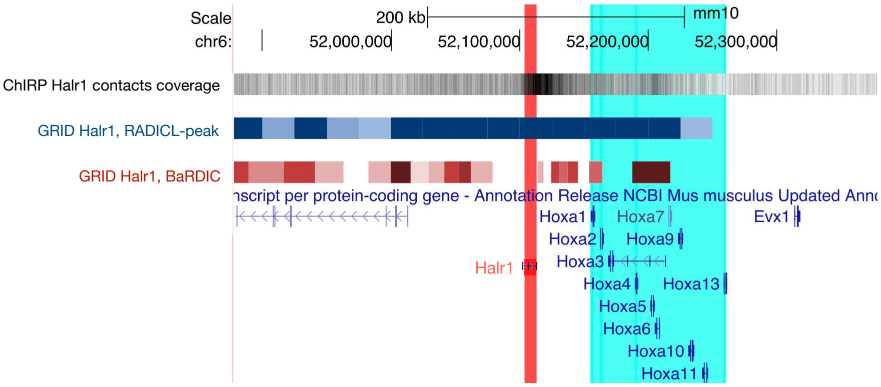


**Supplementary Figure S12:** Representative genome browser view of Halr1 peaks called with BaRDIC and RADICL-peak for GRID ATA data on mESC. Halr1 source gene (red) and HoxA gene cluster (cyan) are highlighted. RADICL peaks are 25 Kb wide, while BaRDIC initial *cis* bin size equals 5 Kb with *cis* factor = 1.12.
