## Supplementary Note for "BaRDIC: robust peak calling for RNA-DNA interaction data"

#### BaRDIC algorithm

The BaRDIC algorithm consists of three steps:

1. Binning: for each RNA, chromosomes are partitioned into non-overlapping genomic intervals (bins), and the number of contacts is calculated within bins.

2. Statistical modeling: for each RNA, parameters of the background model are calculated in every bin. P-values are calculated based on them.

3. Multiple testing correction with Benjamini-Hochberg procedure [1].

To increase the statistical power and speed up the algorithm's performance, by default only RNAs with more than 1000 contacts are selected. Steps 1 and 2 are atomic with regard to individual RNAs, so we describe them for one RNA only.

### 1 Binning

Bin sizes are estimated for each RNA separately. The binning strategy differs for *cis* interactions, which occur on a chromosome harboring the RNA gene, and *trans* interactions – with other chromosomes. To find optimal bin sizes, we adapt RSEG [2] and JAMM [3] approaches that were initially developed for ChIP-seq peak-calling. This approach finds a balance between resolution and sufficient bin coverage.

#### 1.1 Binning in *trans*

In the case of random ligation, we expect that *trans* contacts are distributed uniformly along chromosomes. Therefore, we apply a uniform binning strategy for *trans* interactions. Selection of the optimal bin size for *trans* interactions is done via minimizing the cost function $C$:

$$tran\hat{s}bin = argmin\ C(binsize) = argmin \frac{2 \cdot mean - var}{binsize^2},$$

where *mean* is the average number of contacts in bins, *var* is the variance of the number of contacts in bins, and *binsize* is the corresponding *trans binsize* in nt.

#### 1.2 Binning in *cis*

For binning in *cis* scaling must be taken into account. Also, we assume long-range *cis* interactions are similar to *trans* interactions, analogous to observations in Hi-C data analysis [4]. Since contact densities at different distances can differ by orders of magnitude, we abandoned the uniform binning: chromosomes are partitioned into non-uniform bins of size increasing in a geometric progression from source gene boundaries. *cis* bin size increases until it exceeds the *trans* bin size; all subsequent *cis* bins are uniform and equal to the *trans* bin in size. This strategy achieves:

1. Uniform coverage of bins by contacts, which is required by the bin size optimization procedure.
2. Higher resolution near the source gene and more accurate estimation of *cis* peaks.
3. Similarity of distant *cis* and *trans* bins.

Finally, the size of a *cis* bin with sequence number i from the gene boundaries of a particular RNA is calculated using the formula:

$cis\ binsize(i) = \min(startsize \cdot factor^i, trans\ binsize)$,

where $startsize$ is a tunable user-specified parameter, while $factor$ is optimized by minimizing the cost function similarly to the *trans* bin size selection:

$C(factor) = \frac{2 \cdot mean - var}{factor^2}$.

#### 1.3 Cost function minimization

We found that in our data, the cost function may or may not have a global minimum (Figure 1A, B), so standard function minimization algorithms cannot be applied. Instead, we developed a problem-tailored minimization approach.

We want to find such a bin size that it is not too small (otherwise each bin will have too few contacts, which will reduce statistical power) and not too large (otherwise resolution will be too low). So we start with some small bin size (*startbinsize*), increase it by the same value (*step*) and for each bin size $\{start\ binsize + k \cdot step\}_{k=0}^{\infty}$ we calculate the cost function $\{C_k\}_{k=0}^{\infty}$. Consider the first-order finite difference of the cost function: $\Delta C_k = C_{k+1} - C_k$. Note that it tends to zero as the bin size increases, and then starts to oscillate around it (Figure 1C, D). This behavior means that the rate of decrease of the cost function decreases with increasing bin size up to some point, and then the cost function goes up and down. So, we want to find a point at which the cost function:

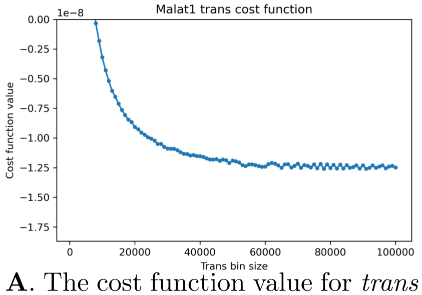


**A**. The cost function value for *trans* bins

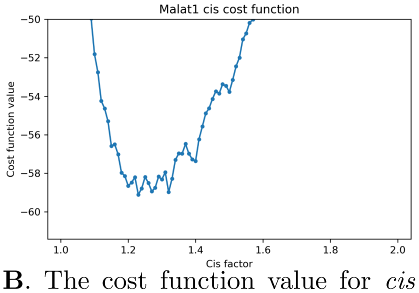


**B**. The cost function value for *cis* factor

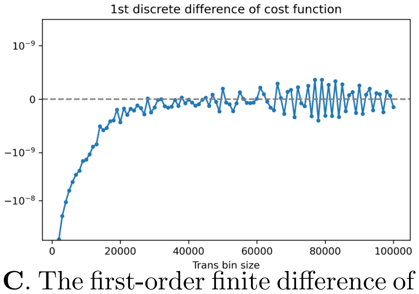


**C**. The first-order finite difference of the cost function for *trans* bins

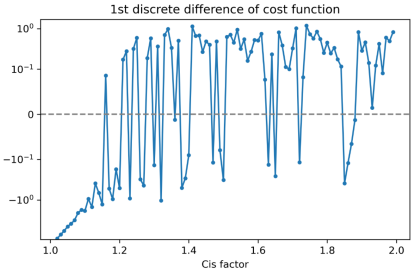


**D**. The first-order finite difference of the cost function for *cis* factor

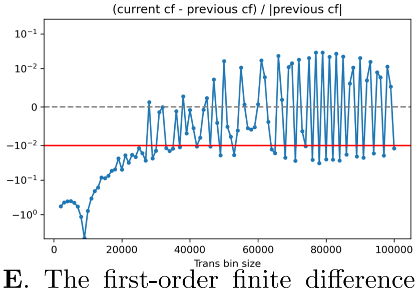


**E**. The first-order finite difference divided by the cost function absolute value for *trans* bins.

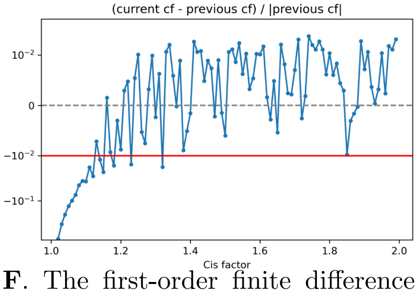


**F**. The first-order finite difference divided by the cost function absolute value for *cis* factor.

**Figure 1:** The optimization process on *trans bin* size and *cis factor* values for MALAT1, GRID-seq data on mESC.

| **Parameter** | ***Trans bin*** | ***Cis factor*** |
|---|---|---|
| start bin size | 10 Kb | 1,1 |
| step | 1 Kb | 0,01 |
| max bin size | 1 Mb | 2 |
| $\delta$ | 0,01 | 0,01 |

**Table 1:** Empirically selected parameters for *cis* and *trans* bins size optimization for all-to-all RNA-DNA interaction data. Set as default parameters of BaRDIC algorithm.

1. Starts decreasing too slowly (there is no global minimum);
2. Or it starts to grow (there is a global minimum).

The second case is defined as $\Delta C_k > 0$, and for the first case we have to introduce some threshold $\delta > 0$ and find a bin size such that $\Delta C_k < \delta$. Since the scale of the cost function will be different for different RNAs and for *trans* and *cis* bins as well, we divide the finite difference by the value of the cost function and take the absolute value of the resulting relative finite difference: $f_k = \frac{\Delta C_k}{|\Delta C_k|}$ (Figure 1 E-F). Then the point of the minimum of the cost function is such a bin size $start\ binsize + kstep$, that corresponds to the smallest $k$ at which one of the three conditions is satisfied:

1. $\Delta C_k > 0$ (the cost function grows);
2. $f_k < \delta$ (the cost function converges);
3. $start\ binsize + k \cdot step > max\ binsize$ (limit, the cost function doesn't converge).

The physical meaning of this optimization procedure is as follows: we choose the smallest bin size that for the (next) larger bin size the cost function either grows or decreases slower than $\delta \cdot 100\%$, or we choose the largest bin size allowed ($max\ binsize$).

For some RNAs, the cost function fluctuates too much for our minimization procedure to work correctly. So the cost function can be smoothed over consecutive bin sizes by taking the average value of the cost function over the previous $w$ steps:

$C_k^{smooth} = \frac{1}{w} \cdot \sum_{h=k-w+1}^{k} C_h$.

By default, no smoothing is performed, i.e. $w = 1$.

The parameters for the optimization procedure were chosen with respect to the resolution of ATA experiments and are presented in Table 1.

### 2 Statistical modeling

#### 2.1 Background model

To model the background distribution of RNA-DNA contacts in bins, we introduce a frequentist model similar to those used in ChIP-seq and Hi-C data analysis [5, 6]. Assuming that contacts arising from random binding are independent, we consider the number of contacts $X_{ij}$ of RNA $i$ in bin $j$ to be binomially distributed:

$$X_{ij} \sim Bin(N_i, p_{ij}),$$

where $N_i$ is the total number of contacts produced by RNA $i$ (except the gene body), $p_{ij}$ is the background probability. Note the number of observed contacts $O_{ij}$ is a realization of the random variable $X_{ij}$. Statistical modeling comes down to inferring the only model parameter $p_{ij}$ from the observed data.

#### 2.2 Inference for *trans* bins

For *trans* bins, only chromatin heterogeneity plays a role. For ATA experiments, we estimate the parameter of the background model by counting mRNA *trans* contacts as proposed in the GRID-peak procedure. The background probability of a single contact of RNA $i$ to appear in the $j$-th *trans* bin is defined as follows:

$$\hat{p} = \hat{p}_j^{bg} = \frac{N_j^{bg}}{N^{bg}},$$

where $N_j^{bg}$ is the number of contacts from the background in a bin $j$, and $N^{bg}$ is the total number of background contacts. For OTA experiments, we use contacts from the input sample similar to ChIP-seq analysis.

#### 2.3 Inference for *cis* bins

To estimate the background probability for *cis* bins, we have to additionally consider scaling. Inspired by Hi-C analysis methods ¡REF Fit-Hi-C¿, we define the corresponding probability as follows:

$$\hat{p_{ij}} = f(d_{ij}) \cdot \hat{p_j^{bg}},$$

where $d_{ij}$ is the distance between the midpoint of bin $j$ and the closest boundary of RNA $i$ source gene, $f(d_{ij})$ – is the scaling factor. Assuming $O_{ij} \approx EX_{ij} = N_i\hat{p_{ij}}$, we can estimate $f(d_{ij})$ as follows: $\hat{f}(d_{ij}) = \frac{O_{ij}}{N_i \cdot \hat{p_j^{bg}}}$.

Since the bins on the 5'- and 3'-sides of a source gene are at the same distance from the gene, there are two values of $\hat{f}$ for each $d_{ij}$; however, there should be only one value of $f$. We calculate $f$ using a smoothing B-spline of degree 3 in double logarithmic coordinates. It turns out that for individual RNAs from a single chromosome, $\hat{f}(d_{ij})$ may differ by 2-3 orders of magnitude in (Figure 2), probably due to the sparsity of the data. Accordingly, the spline is calculated separately for each RNA.

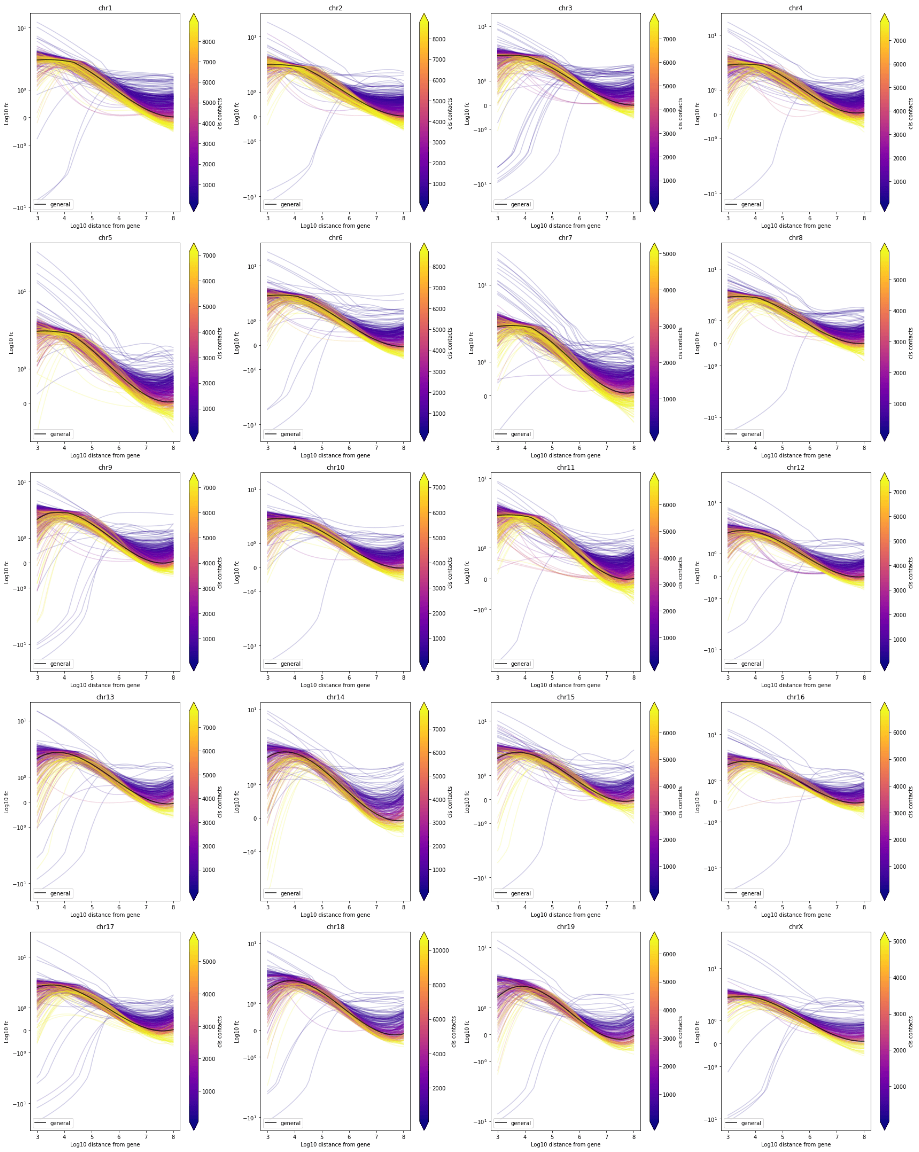


**Figure 2:** Scaling splines of individual RNAs in all-to-all GRID data on mESC cell line.

To exclude potential *cis* peaks from the background model, we estimate scaling factors in 2 steps similar to Fit-Hi-C [5] and HiC-DC [7]. In the first step, we compute spline-1 over all bins $j$ for each RNA $i$, then determine p-values by right-sided binomial test for each *cis* bin and remove those bins with $pvalue = \frac{1}{M_i^{cis}}$, where $M_i^{cis}$ is the number of non-zero *cis* bins for RNA $i$. If $\frac{1}{M_i^{cis}}$ is greater than the threshold value 0.05, the latter is used. This filtering excludes likely specific binding sites. We then compute spline-2 in the same manner using only the remaining bins. When computing both splines, we remove all bins $j$ for which $O_{ij} = 0$ and/or $N_j^{bg} = 0$ to avoid spline distortion due to data sparsity.

The final scaling factor for each RNA $i$ and each bin $j$ is calculated as the geometric mean of three scaling factors: spline-2 values for the beginning of the bin, the middle of the bin, and the end of the bin. "1" is added to each relative coordinate to avoid zeros when taking logarithms.

#### 2.4 Treating zero values

In both *cis* and *trans* inference, except spline fitting, the background coverage value $N_j^{bg} = 0$ is treated as missing and imputed with a very small positive value. We define it as the average base-pair coverage of the genome by background contacts, multiplied by the length of bin $j$ and by a reducing imputation factor (0.01 by default).

#### 2.5 Parameter renormalization

To hold $\sum_j \hat{p}_{ij} = 1$, we renormalize probability estimates for each RNA $i$. For this purpose, it is easiest to partition the probability space of the whole genome into two subspaces: the source RNA chromosome and the other chromosomes, so that the sum of the background probabilities for *cis* bins equals the fraction of *cis* contacts of this RNA (similarly for *trans* bins). For this purpose, let us introduce some more notations:

| **Notation** | **Meaning** |
|---|---|
| $C(i)$ | The source chromosome of RNA $i$ |
| $C(j)$ | The chromosome of bin $j$ |
| $N_i^{cis} \equiv \sum_{j:C(i)=C(j)} O_{ij}$ | The number of *cis* contacts of RNA $i$ |
| $\hat{p}_j^{bg^{cis}} \equiv \frac{N_j^{bg}}{\sum_{j:C(i)=C(j)} N_j^{bg}}$ | The frequency of background contacts in *cis* for RNA $i$ |
| $\hat{p}_j^{bg^{trans}} \equiv \frac{N_j^{bg}}{\sum_{j:C(i)\neq C(j)} N_j^{bg}}$ | The frequency of background contacts in *trans* for RNA $i$ |

Then, renormalized estimated background probabilities $\hat{p}_{ij}$ are calculated as follows:

| Type of bin | Background probability | Sum of background probabilities |
|---|---|---|
| *trans* | $\hat{p}_{ij}^{trans} = \hat{p}_j^{bg^{trans}} \cdot \frac{N_i - N_i^{cis}}{N_i}$ | $\sum_j \hat{p}_{ij}^{trans} = \frac{N_i - N_i^{cis}}{N_i}$ |
| *cis* | $\hat{p}_{ij}^{cis} = \frac{\hat{f}_i(d_{ij}) \cdot \hat{p}^{bg^{cis}}}{\sum_{j:C(i)=C(j)} \cdot \hat{f}_i(d_{ij}) \cdot \hat{p}^{bg^{cis}}} \cdot \frac{N_i^{cis}}{N_i}$ | $\sum_j \hat{p}_{ij}^{cis} = \frac{N_i^{cis}}{N_i}$ |

These background probabilities are the final parameter values ($\hat{p}_{ij}$) of the background binomial model.

##### 2.6 P-value calculation

We expect that a specific binding event results in a high enrichment of contacts relative to the background model. The resulting $p - value$ is then computed with the right-sided binomial test using estimated background parameters:

$$pvalue(O_{ij}|X_{ij}) = P_{Bin}(X_{ij} \geq O_{ij}|N_i, \hat{p}_{ij}) = \sum_{k=O_{ij}}^{N_i} \binom{N_i}{k} \cdot \hat{p}_{ij}^k \cdot (1 - \hat{p}_{ij})^{N_i - k}$$

#### 3 Multiple testing correction

*P-values* from non-zero bins are subjected to multiple testing correction using the Benjamini-Hochberg procedure, simultaneously for all RNAs. This procedure controls FDR on the global level, for all peaks of all RNAs in an ATA dataset. So the same FDR level cannot be propagated to peaks of individual RNAs based on this procedure.

#### 4 Implementation

##### 4.1 Software

The algorithm is implemented as a package for Python 3 [8] with a command line interface and is available at https://github.com/dmitrymyl/BaRDIC. It uses several packages for scientific computing (see the table below). Since binning and statistical evaluation are performed for each RNA separately, these steps are parallelized.

| Package name | Version | Usage | Reference |
|---|---|---|---|
| numpy | 1.24.4 | Vector data operations | [9] |
| pandas | 2.0.3 | Table data operations | [10] |
| scipy | 1.10.1 | Splines and binomial tests | [11] |
| statsmodels | 0.14.0 | FDR correction | [12] |
| bioframe | 0.4.1 | Operations on genomic intervals | [13] |
| h5py | 3.9.0 | Access to HDF5 storage | [14] |
| tqdm | 4.65.0 | Progress bars and process-based parallelization via concurrent.futures | [15] |

#### 4.2 Data storage

To organize the data storage and speed up the data access, we developed two HDF5-based data formats [14].

##### 4.2.1 dnah5 file format

This file format is a binary representation of DNA parts of contacts grouped by individual RNAs with optimized bin sizes. The layout of the dnah5:

```
/
- chrom_sizes/
    - chrom | O
    - size | int64
- dna_parts/
    - rnaN/
        - chromN/
            - start | int64
            - end | int64
```

File-level attributes:

| Attribute | Type | Description |
|---|---|---|
| are_binsizes_selected | bool | Whether bin sizes are selected for each RNA and corresponding data is recorded in the file |
| version | str | dnah5 schema version.<br>Currently, only “1” is available |

RNA-level attributes contain coordinates of the source gene, contact statistics, and binning parameters:

| Field | Type | Description |
|---|---|---|
| chrom | O | RNA gene chromosome |
| start | int64 | RNA gene start |
| end | int64 | RNA gene end |
| total_contacts | int64 | Total number of RNA contacts |
| genic_contacts | int64 | Number of RNA contacts inside its gene |
| *cis*_contacts | int64 | Number of *cis* RNA contacts: on the RNA's origin chromosome but outside RNA gene |
| *trans*_contacts | int64 | Number of *trans* RNA contacts: on all chromosomes except for RNA's origin one |
| eligible | bool | Whether this RNA has enough contacts for further processing |
| *cis*_factor | float64 | *cis* factor value for binning |
| *cis*_start | int64 | Initial *cis* bin size |
| *trans*_bin_size | int64 | *trans* bin size |

##### 4.2.2 rdc file format

This file format holds binned RNA-DNA contacts and corresponding values as well as the binned background track. Bin tables are organized for each RNA separately. The layout of the rdc:

```
/
- chrom_sizes/
    - chrom | O
    - size | int64
- background
    - chrN
        - start | int64
        - end | int64
        - count | int64
- pixels
    - rnaN
        - chrN
            - start | int64
            - end | int64
            - signal_count | float64
            - bg_count | float64
            - raw_bg_prob | float64
            - scaling_factor | float64
            - bg_prob | float64
```

```
- impute | bool
- signal_prob | float64
- fc | float64
- pvalue | float64
- qvalue | float64
```

File-level attributes:

| Field | Type | Description |
|---|---|---|
| is_scaling_fitted | bool | Whether scaling is estimated and RNAs background levels of interactions are rescaled |
| are_peaks_estimated | bool | Whether p-values and q-values for peaks are estimated |
| version | str | rdc schema version. Currently, only “1” is available |

RNA-level attributes contain source gene coordinates, contact statistics, binning, and spline parameters:

| Field | Type | Description |
|---|---|---|
| chrom | O | RNA gene chromosome |
| start | int64 | RNA gene start |
| end | int64 | RNA gene end |
| total_contacts | int64 | Total number of RNA contacts |
| genic_contacts | int64 | Number of RNA contacts inside its gene |
| *cis*_contacts | int64 | Number of *cis* RNA contacts: on the RNA's origin chromosome but outside RNA gene |
| *trans*_contacts | int64 | Number of *trans* RNA contacts: on all chromosomes except for RNA's origin one |
| eligible | bool | Whether this RNA has enough contacts for further processing |
| *cis*_factor | float64 | *cis* factor value for binning |
| *cis*_start | int64 | Initial *cis* bin size |
| *trans*_bin_size | int64 | *trans* bin size |
| scaling_spline_t | Float array | A vector of knots of a scaling B-spline |
| scaling_spline_c | Float array | A vector of scaling B-spline coefficients |
| scaling_spline_k | int64 | A degree of a scaling B-spline |

[10] Jeff Reback, Wes McKinney, jbrockmendel, Joris Van den Bossche, Tom Augspurger, Phillip Cloud, gfyoung, Sinhrks, Adam Klein, Matthew Roeschke, Jeff Tratner, Chang She, William Ayd, Simon Hawkins, Terji

Petersen, Jeremy Schendel, Andy Hayden, Marc Garcia, Vytautas Jancauskas, MomIsBestFriend, Pietro Battiston, Skipper Seabold, chris b1, h vetinari, Stephan Hoyer, Wouter Overmeire, alimcmaster1, Mortada Mehyar, Christopher Whelan, and Thomas Kluyver. pandas-dev/pandas: Pandas 1.0.0, January 2020.

[11] Pauli Virtanen, Ralf Gommers, Travis E. Oliphant, Matt Haberland, Tyler Reddy, David Cournapeau, Evgeni Burovski, Pearu Peterson, Warren Weckesser, Jonathan Bright, Stéfan J. van der Walt, Matthew Brett, Joshua Wilson, K. Jarrod Millman, Nikolay Mayorov, Andrew R. J. Nelson, Eric Jones, Robert Kern, Eric Larson, C J Carey, İlhan Polat, Yu Feng, Eric W. Moore, Jake VanderPlas, Denis Laxalde, Josef Perktold, Robert Cimrman, Ian Henriksen, E. A. Quintero, Charles R. Harris, Anne M. Archibald, Antônio H. Ribeiro, Fabian Pedregosa, Paul van Mulbregt, Aditya Vijaykumar, Alessandro Pietro Bardelli, Alex Rothberg, Andreas Hilboll, Andreas Kloeckner, Anthony Scopatz, Antony Lee, Ariel Rokem, C. Nathan Woods, Chad Fulton, Charles Masson, Christian Häggström, Clark Fitzgerald, David A. Nicholson, David R. Hagen, Dmitrii V. Pasechnik, Emanuele Olivetti, Eric Martin, Eric Wieser, Fabrice Silva, Felix Lenders, Florian Wilhelm, G. Young, Gavin A. Price, Gert-Ludwig Ingold, Gregory E. Allen, Gregory R. Lee, Hervé Audren, Irvin Probst, Jörg P. Dietrich, Jacob Silterra, James T Webber, Janko Slavič, Joel Nothman, Johannes Buchner, Johannes Kulick, Johannes L. Schönberger, José Vinícius de Miranda Cardoso, Joscha Reimer, Joseph Harrington, Juan Luis Cano Rodríguez, Juan Nunez-Iglesias, Justin Kuczynski, Kevin Tritz, Martin Thoma, Matthew Newville, Matthias Kümmerer, Maximilian Bolingbroke, Michael Tartre, Mikhail Pak, Nathaniel J. Smith, Nikolai Nowaczyk, Nikolay Shebanov, Oleksandr Pavlyk, Per A. Brodtkorb, Perry Lee, Robert T. McGibbon, Roman Feldbauer, Sam Lewis, Sam Tygier, Scott Sievert, Sebastiano Vigna, Stefan Peterson, Surhud More, Tadeusz Pudlik, Takuya Oshima, Thomas J. Pingel, Thomas P. Robitaille, Thomas Spura, Thouis R. Jones, Tim Cera, Tim Leslie, Tiziano Zito, Tom Krauss, Utkarsh Upadhyay, Yaroslav O. Halchenko, and Yoshiki Vázquez-Baeza and. SciPy 1.0: fundamental algorithms for scientific computing in python. *Nature Methods*, 17(3):261–272, February 2020.

[12] Skipper Seabold and Josef Perktold. Statsmodels: Econometric and statistical modeling with python. In *Proceedings of the Python in Science Conference*. SciPy, 2010.

[13] Open2C, Nezar Abdennur, Geoffrey Fudenberg, Ilya Flyamer, Aleksandra A. Galitsyna, Anton Goloborodko, Maxim Imakaev, and Sergey V. Venev. Bioframe: Operations on genomic intervals in pandas dataframes. February 2022.

[14] The HDF Group. Hierarchical data format version 5, 2000-2010.

[15] Casper da Costa-Luis, Stephen Karl Larroque, Kyle Altendorf, Hadrien Mary, richardsheridan, Mikhail Korobov, Noam Yorav-Raphael, Ivan Ivanov, Marcel Bargull, Nishant Rodrigues, Guangshuo Chen, Antony Lee, Charles Newey, CrazyPython, JC, Martin Zugnoni, Matthew D. Pagel, mjstevens777, Mikhail Dektyarev, Alex Rothberg, Alexander Plavin, Fabian Dill, FichteFoll, Gregor Sturm, HeoHeo, Hugo van Kemenade, Jack McCracken, MapleCCC, Max Nordlund, and Mike Boyle. tqdm: A fast, Extensible Progress Bar for Python and CLI, August 2023.
